# Better null models for assessing predictive accuracy of disease models

**DOI:** 10.1101/2021.07.26.453866

**Authors:** Alexander C. Keyel, A. Marm Kilpatrick

**Author notes:** Contact information: ACK; 120 New Scotland Road, Albany, NY 12208., AMK. Statement of Authorship: ACK and AMK designed the study, drafted and revised the manuscript, and performed the analyses. Data Accessibility Statement: No new data were used in this study. Data used in this manuscript are available from the Centers for Disease Control.

## Abstract

Null models provide a critical baseline for the evaluation of predictive disease models. Many studies consider only the grand mean null model (i.e. R^2^) when evaluating the predictive ability of a model, which is often misleading in conveying the predictive power of a model. We evaluated ten null models for human cases of West Nile virus (WNV), a zoonotic mosquito-borne disease introduced to the United States in 1999. The Negative Binomial, Historical (i.e. using previous cases to predict future cases) and Always Absent null models were the strongest overall, and the majority of null models significantly outperformed the grand mean. Somewhat surprisingly, the length of the training timeseries did not strongly affect the performance of most null models. We argue that a combination of null models is needed to evaluate the forecasting performance of predictive models and the grand mean is the lowest bar.

## Introduction

Forecasting infectious disease dynamics is a key challenge for the 21^st^ century (Johansson *et al*. 2019). Climate and land use change, combined with the introduction of pathogens to new regions, has created an urgent need for predicting future disease threats (Kilpatrick & Randolph 2012). Large data sets and new modeling and statistical techniques have opened up possibilities for ecological forecasting (Dietze 2017). A key step in the evaluation of predictive models is assessing their improvement over null models. The use of null models to provide a baseline in the absence of specific mechanisms has a long history in ecology (Gotelli & Graves 1996). Such baselines are important, as in some cases, predictive models may appear to be informative, but may be no better than a simple and uninformative null model (Olden *et al*. 2002; Beale *et al*. 2008). For example, when dealing with rare events, if a predictive model is outperformed by a null model that predicts the event to never occur, it is not providing much useful information about the process being studied (Olden *et al*. 2002).

West Nile Virus (WNV) is an excellent system in which to examine null models in a probabilistic context. WNV is a flavivirus that cycles between mosquito and avian populations (Work *et al*. 1955; Komar *et al*. 2003; Kilpatrick 2011). WNV was introduced to the United States (US) in 1999 (Lanciotti *et al*. 1999) and rapidly spread to the conterminous US and throughout the Americas (Kramer *et al*. 2019). As a nationally notifiable disease in the US, long-term data sets (>20 years) exist on human cases (CDC 2019b). Many models have been built for predicting WNV risk (Barker 2019). Most studies of WNV, and many other pathogens, have included only a very simplistic null model (e.g. R^2^, which uses the grand mean of the training data) for assessment of model accuracy.

Our aim was to examine a range of null models (Box 1) to provide guidance on null model selection and performance in disease forecasting for locations with frequent (≥50% of years with disease) and infrequent cases (disease present, but <50% of years). We tested 10 null models using the number of WNV cases in each county in the US in each year in a probabilistic framework. Where cases were frequent and timeseries were long, we hypothesized the Negative Binomial model would perform the best due to its ability to model count distributions with a variable rate parameter. Where cases were infrequent and time series were short, we predicted that no models would significantly outperform the Always Absent mode.

## Material and methods

### Data set

We compared the accuracy of 10 null models using the CDC Neuroinvasive WNV Case records (Source: ArboNET, Arboviral Diseases Branch, Centers for Disease Control and Prevention, Fort Collins, Colorado; contact the CDC for data access). This is a national data set of the number of WNV neuroinvasive disease cases in each county in each year from 3108 counties in the conterminous United States (US) from 2000 – 2021. We used WNV neuroinvasive cases, because there is less variability across different states in detection of these cases compared to WNV fever cases.

We divided the data set into two groups: 159 counties that have had 11 or more years with WNV (frequent WNV set), and 1880 counties that had 1 – 10 years with WNV (infrequent WNV set). The 1069 counties that never had WNV cases were excluded to avoid zero-inflation. The first year a state reported a case of WNV (per (Kilpatrick 2011)) was used as the first year of training data for all counties within that state. As a result, the number of counties included in the analysis increased over time (S1 Table). Model predictions were made using at least 4 years of training data. We used the Continuous Ranked Probability Score, a probabilistic scoring approach that can evaluate a distribution of predicted outcomes (Box 2). Population data for each year for the incidence-based null models came from the United States Census Bureau (US Census Bureau 2017, 2019). Population data from 2019 were used for 2020 and 2021 as well, due to missing data for these years.

We also tested whether our model results were sensitive to the length of time series for selected models. Model years were selected at random (without replacement) to use as training data to predict a randomly selected focal year. This allowed us to disentangle length of time series from the specific order of observation of results. However, models that required a temporal structure were excluded from this analysis (i.e. Prior Year and Autoregressive). The Incidence and Pooled Incidence models were also excluded from this analysis as they used the prior-year’s population for converting incidence to case counts. Only data from 2005 and later were used in this analysis to ensure that WNV had already been established in all counties.

### Null Models

The ten null models are described in Box 1. Note that using case counts versus incidence does not make a large difference to the outcome when stratifying by county, because population is relatively consistent from one year to another. However, the choice of incidence or case counts as the model basis does lead to different outcomes when pooling across counties with different population sizes.

### Scoring Method

We used the Continuous Ranked Probability Score (CRPS), which is a proper scoring method (Jordan *et al*. 2019; Bracher *et al*. 2021) We chose the CRPS rather than the Logarithmic Score because the former scores forecasts based on the distance from each predicted probability to the observation, whereas the latter only scores whether an observation is within a bin or outside of a bin, with no consideration of how far outside the bin the prediction was (Matheson & Winkler 1976; Hersbach 2000; Wilks 2011). As the CRPS scores are based on distance from the observed value, the models above were allowed to predict fractional cases of WNV (e.g. if the mean number of cases was 2.5 cases, that would be used as the prediction rather than rounding up or down to the nearest whole number). For null models that required sampling from a probabilistic distribution, we used 100 random draws. Data analyses were performed in R (R Core Team 2017). Code for running the null models is available via the probnulls package on GitHub (www.github.com/akeyel/probnulls/R/NullModels.R).

## Results

The Mean Value null model (R^2^) that is frequently used as the baseline for prediction accuracy was among the weakest of the null models (Fig. 1). It performed worse than 5 null models (significantly worse than 4; Fig. 1) for frequent WNV counties and worse than 6 null models (significantly worse than 5) for infrequent WNV counties (Fig. 1). In contrast, the Negative Binomial null model was significantly better than other null models for predicting neuroinvasive cases of WNV in the frequent WNV analysis (Fig. 1a, paired t-tests using a Holm correction for multiple comparisons). The Negative Binomial was also significantly better than eight other null models (all except the Always Absent model which was equally accurate) in the infrequent WNV analysis (Fig. 1b). The Negative Binomial was the top model in 8 individual years (out of 18) for the frequent WNV analysis, and in 9 of 18 years for the infrequent WNV analysis (Table 1). The Historical Null model also performed very well in both frequent and infrequent WNV analyses across all years (Fig 1), and outperformed all other models in 5 individual years for frequent WNV counties (Table 1). Finally, in the infrequent WNV analysis, the Always Absent model was tied for the best model for all years combined, and outperformed seven other models (Fig 1b) and was the best model for 8 individual years (Table 1).

**Table 1.**
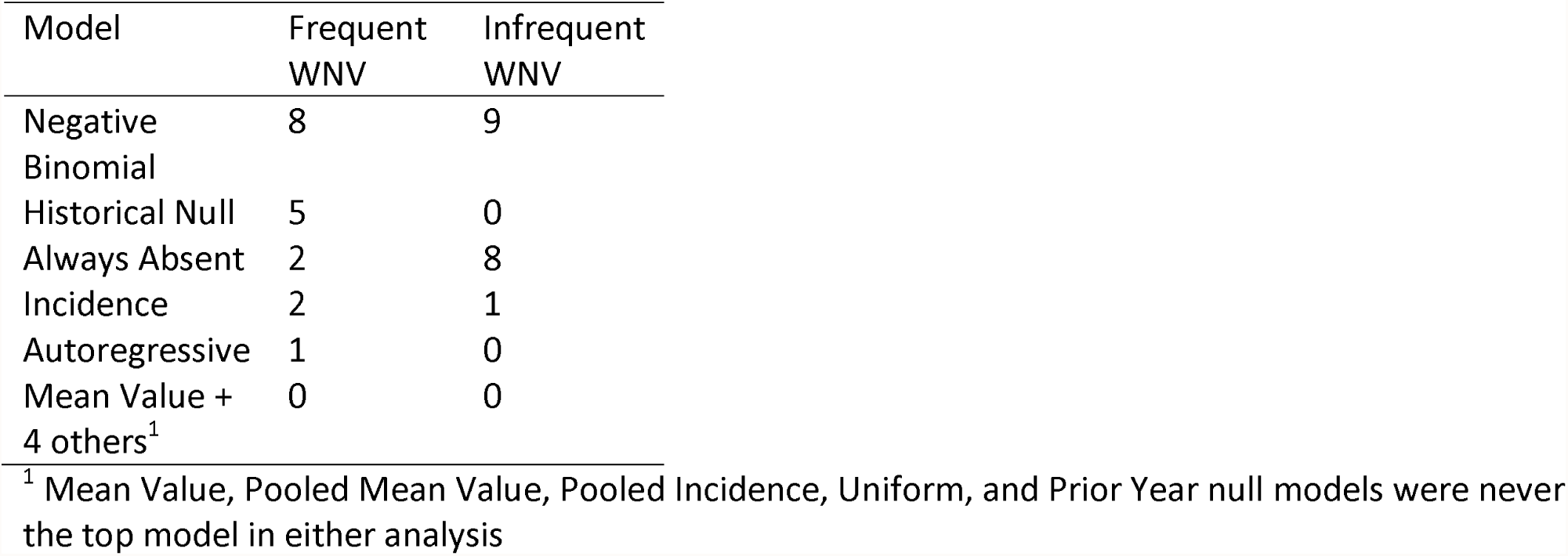
Frequency of each model having the lowest CRPS score for a year in counties where WNV is frequent (>50% of time series) or infrequent (present < 50% of the time series).

**Fig. 1.**
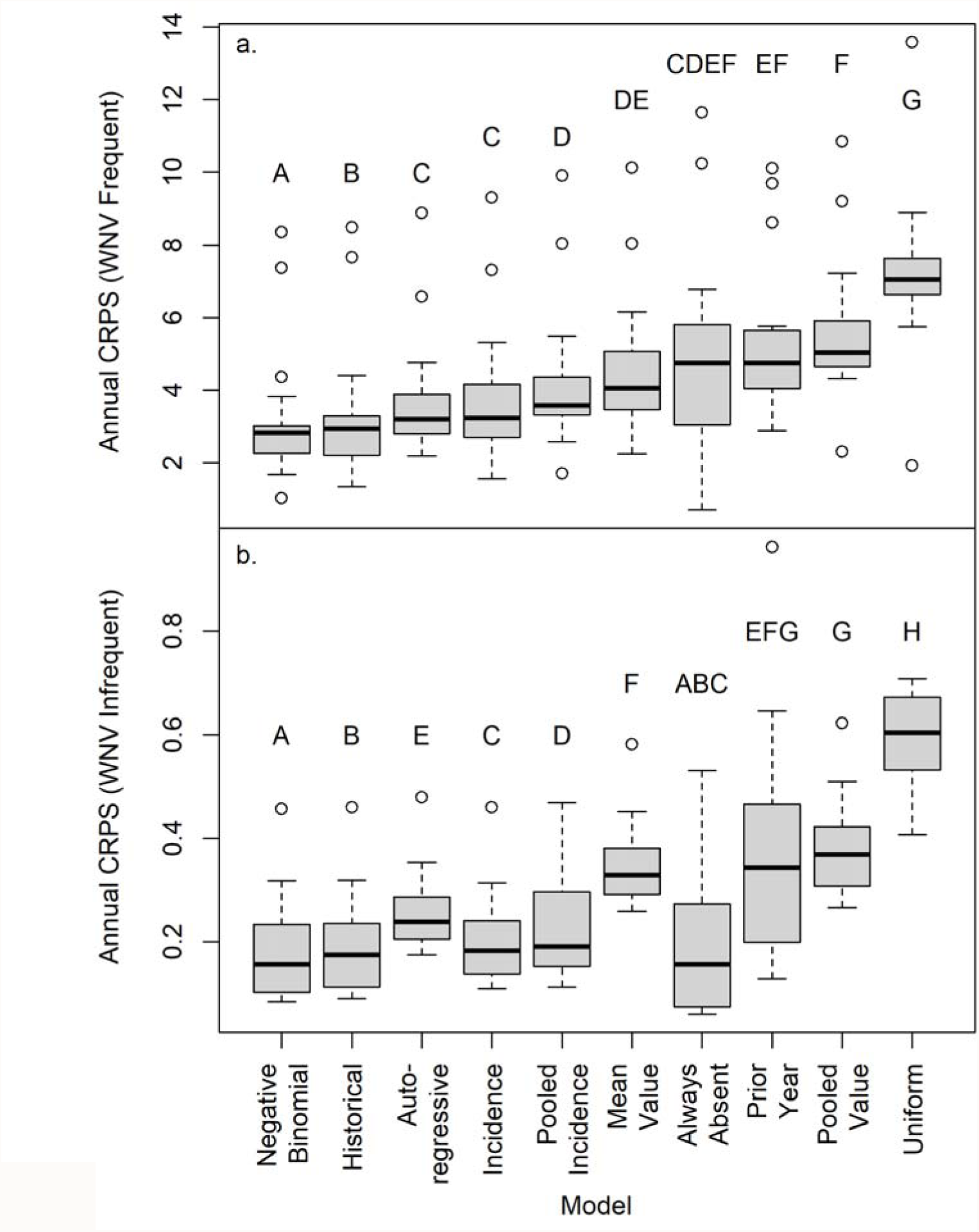
Continuous Ranked Probability Scores (CRPS) for 2004 – 2021 for 10 null models for a) counties with at least 50% of the time series with WNV (“Frequent”) and b) for counties with at least one case, but cases < 50% of the time series (“Infrequent”). The mean CRPS scores was calculated across all counties for each model and year, and the plot shows the median value of these mean annual CRPS scores by model, with the box showing the 25% and 75% quartiles, whiskers corresponding to +/- 1.5 times the Interquartile range, and circles corresponding to values outliers outside this range.

The length of the training time series had only weak effects on null model performance (Fig. 2). For infrequent WNV counties there was no support for an improvement in model score with the length of the training time series (Table S2). For frequent WNV counties, model score improved with the length of the training time series for four of the six models examined, but the effect was small relative to the difference between most models (Table S3). For the two remaining models, increasing the length of the training time series had no effect on model score or actually made the score worse (Table S3).

**Fig. 2.**
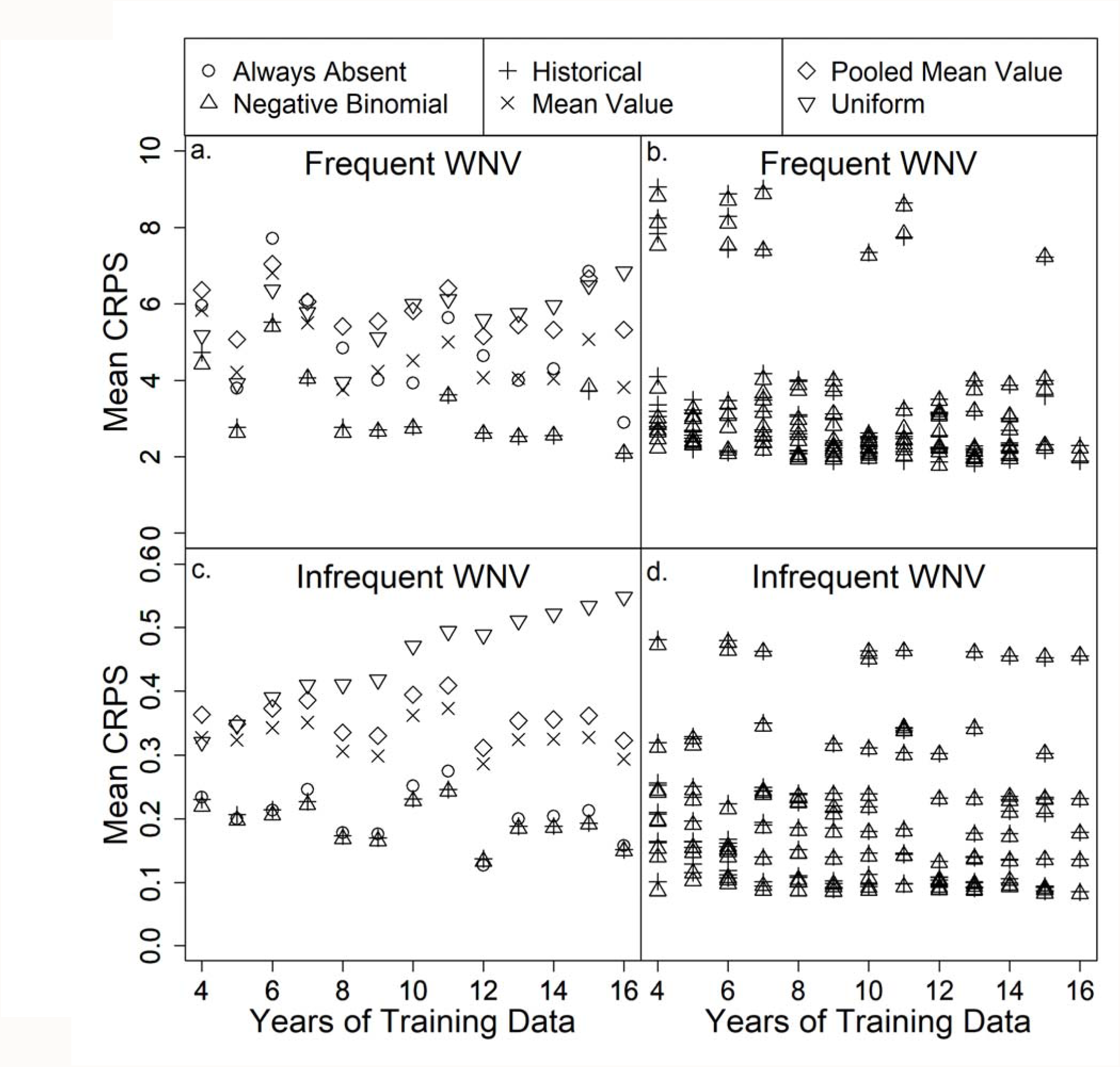
The Negative Binomial and Historical were generally the top two models, independent of length of time series used to train the models for both a) the counties with frequent WNV and c) the counties with infrequent WNV cases. Individual training replicates showed a range of variation in CRPS score (b, d) depending on which replicates were selected (only Negative Binomial and Historical null models shown). Training years were randomly selected from the entire time series, and a random focal year was selected for evaluation. Only a subset of null models was evaluated over time.

## Discussion

At least five null models significantly outperformed a county-based grand mean and many did far better (Figs. 1, 2). A grand mean calculated for the entire US (Pooled Mean model) performed even worse. Thus, when evaluating the performance of new statistical or mechanistic models of disease incidence, there are far better null models than the grand mean (i.e. R^2^). These null models can be easily calculated for time-series data (e.g., using the probnulls package from GitHub in R), and our results suggest that the length of time series was not critical for developing a robust null model across a range of 4-16 years. The Negative Binomial and Historical nulls were the strongest null models overall, with the Always Absent null performing well where disease cases were infrequent. The strong performance of the Always Absent null in regions where WNV was infrequent (statistically tied with Negative Binomial, Fig. 1; top model in 8 of 18 years, Table 1) is a reminder that basic accuracy statistics for rare events can appear high.

The structure and scale of the underlying data may affect the performance of the different null models. The WNV data set here does not have a clear temporal trend. A strong temporal trend would likely have changed which model performed the best. Specifically, null models that use the recent past to predict future cases (e.g. autoregressive models) would perform much better. Seasonal patterns, as examined in recent dengue forecasts (Johansson *et al*. 2019), could also affect which null model performs best. Future work could explore the performance of different models under different magnitudes of temporal trend and stochastic variation. Additionally, county-annual scales may be more relevant to academic study than to vector control and public health responses (Keyel *et al*. 2021). Research on null model performance is needed at finer spatial and temporal scales.

### Conclusion

We strongly recommend the inclusion of multiple null models when testing predictive models of vector-borne diseases. A grand mean calculated from the training data set is an inadequate null model given the suite of probabilistic alternatives available. The Negative Binomial and Historical nulls performed especially well for WNV and simple autoregressive models performed moderately well and would likely perform even better for data with temporal trends. Negative Binomial and Historical null models performed well both when WNV cases were frequent and when they were infrequent, and their performance did strongly not depend on the length of the training time series. Researchers proposing mechanistic models should determine if their models are an improvement over a simple statistical description of historical patterns.

## Acknowledgements

We thank L. F. Chaves for constructive discussion. This publication was supported by cooperative agreement 1U01CK000509-01, funded by the Centers for Disease Control and Prevention and by the National Institutes of Health grant R01AI168097 and National Science Foundation grants DEB 1911853, DEB-1717498 and CNH-1115069. Its contents are solely the responsibility of the authors and do not necessarily represent the official views of the Centers for Disease Control and Prevention or the Department of Health and Human Services.

---

**Box 1. Meet the 10 Null Models**

**Always Absent Null** (non-probabilistic): WNV is always predicted to be absent (i.e., the model always predicts 0 cases or incidence).

**Pooled Mean Value Null^1^** (non-probabilistic): WNV is predicted to have its mean value of cases.

**Mean Value Null** (non-probabilistic): As in the Pooled Mean Value Null, but stratified by county.

**Prior Year Null** (non-probabilistic): The results from the prior year are used to predict the current year (Smith *et al*. 2020).

**Historical Null:** Prior observations for the location are sampled with replacement, creating a probabilistic distribution of future cases.

**Pooled Incidence Null:** The number of cases were predicted by sampling a binomial distribution, using incidence as the probability, and the county’s population as the size parameter. Incidence was calculated for the entire region of interest (i.e. the United States).

**Incidence Null:** As in the Pooled Incidence Null, but incidence is calculated stratified by county, allowing the model to capture local hotspots of disease.

**Negative Binomial Null^2^**: We estimated the mean and dispersion parameters of a negative binomial distribution using the human case counts in each year from each county Random draws for each county were then drawn using the estimated mean and dispersion parameter.

**Autoregressive AR1 Null^3^:** An autoregressive model was fit. The model produced a mean estimate for the next time step and a normally-distributed error around that mean. We randomly sampled the normal distribution around the mean. All values less than zero were assigned to 0, as negative human cases are not possible.

**Uniform Null:** Random draws were taken from a uniform distribution of cases bounded by 0 to the maximum number of cases observed in a county.

**Box 2. A quick introduction to probabilistic models**

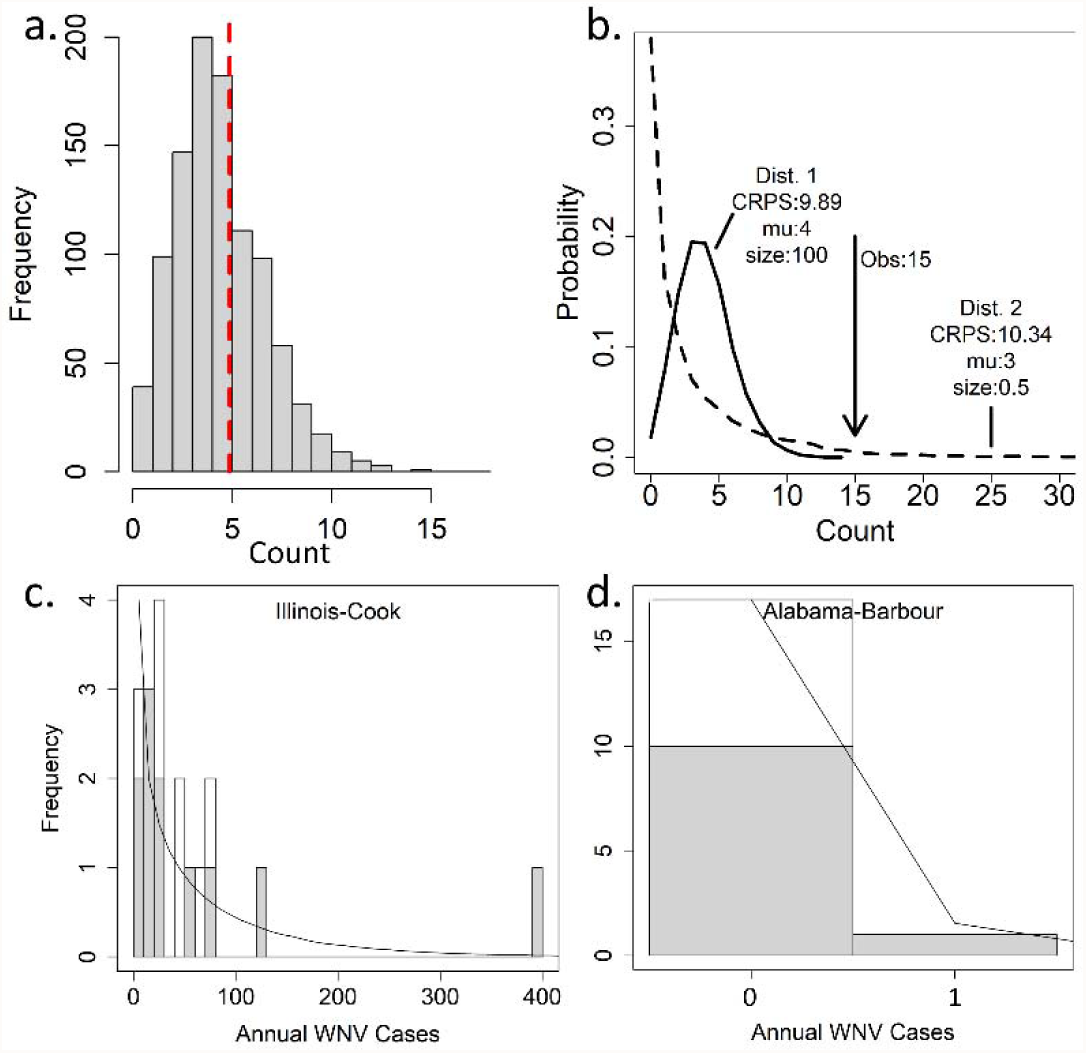

A probabilistic forecast can convey substantially more information than a point forecast. Here, (a) a dashed line indicates the point estimate, while the histogram of 1000 samples from a negative binomial distribution with μ = 5 (mean) and size = 100 (dispersion parameter) show the expected range of possible outcomes. (b) Note that because the CRPS score is based on distance between the observed value and the entire distribution of the prediction, it can give non-intuitive results where a distribution that includes the observed value (Dist. 2) can score worse than a distribution that assigns the observation a zero or near-zero probability (Dist. 1). (c) Negative binomial fits for a county with a high number of annual neuroinvasive West Nile virus (WNV) cases and (d) a county with very few WNV cases. Shading shows data used to train the models (2002 - 2012) and unshaded shows testing data used to evaluate the models (2013 - 2019). The contribution of each year to the overall CRPS score is given in the table below for the two locations for three null models: Always Absent, Historical, and Negative Binomial. The CRPS value for the Always Absent Null is also the number of WNV cases for that year because the CRPS becomes the mean absolute error when applied to a point estimate and the Always Absent Null always predicts 0 cases. Lower CRPS scores indicate better model performance, with 0 being a perfect prediction. Here, Historical performed the best on average for Cook, IL, while Always Absent performed the best on average for Barbour, AL. Visually, the Negative Binomial fit for Cook County appears poor, due to over-estimation in the lowest case bin, and under-estimation of most of the remaining bins.

| Year | Cook, IL |  |  | Barbour, AL |  |  |
| --- | --- | --- | --- | --- | --- | --- |
|  | Always Absent | Historical | Negative Binomial | Always Absent | Historical<sup>1</sup> | Negative Binomial<sup>1</sup> |
| 2013 | 46 | 15.9 | 16.1 | 0 | 0.008 | 0.008 |
| 2014 | 23 | 8.3 | 14.6 | 0 | 0.008 | 0.008 |
| 2015 | 21 | 8.2 | 14.8 | 0 | 0.008 | 0.008 |
| 2016 | 65 | 24.9 | 21.9 | 0 | 0.008 | 0.008 |
| 2017 | 41 | 14.1 | 15.3 | 0 | 0.008 | 0.008 |
| 2018 | 76 | 31.5 | 26.7 | 0 | 0.008 | 0.008 |
| 2019 | 9 | 11.3 | 19.4 | 0 | 0.008 | 0.008 |
| Mean | 40.1 | 16.3 | 18.4 | 0 | 0.008 | 0.008 |

<sup>1</sup> By chance, these two models predicted the same distribution of zero and one case occurrences. Different random draws would cause these scores to vary slightly.

**S1 Table.** Number of counties used for the null model analysis by prediction year for 2004 – 2021. After 2008, the sample size remained constant for all following years as all states had the minimum 4 years of training data with WNV by that point.

| Year | N (frequent<br>WNV) | N (infrequent<br>WNV) |
| --- | --- | --- |
| 2004 | 25 | 213 |
| 2005 | 79 | 1042 |
| 2006 | 154 | 1831 |
| 2007 | 159 | 1865 |
| 2008 – 2021 | 159 | 1880 |

**Table S2.** **Analysis of the length of the training time series on the CRPS score for six null models for infrequent WNV counties. An additive model of null model and time series length fit better than an interactive model (ΔAIC = 50)**.

| Predictor | Estimate | SE | t value | P-value |
| --- | --- | --- | --- | --- |
| Intercept (Neg Binomial) | 0.18 | 0.015 | 12.18 | <2e-16 |
| Length of time series | 0.00072 | 0.0011 | 0.68 | 0.50 |
| Always absent | 0.014 | 0.014 | 1.01 | 0.31 |
| Historical | 0.0047 | 0.014 | 0.34 | 0.73 |
| Mean | 0.13 | 0.014 | 9.82 | <2e-16 |
| Pooled mean | 0.17 | 0.014 | 12.09 | <2e-16 |
| Uniform | 0.26 | 0.014 | 18.94 | <2e-16 |

**Table S3.** **Analysis of the length of the training time series on the CRPS score for six null models for frequent WNV counties. A model with an interaction between model and time series length fit better than an additive model (ΔAIC = 21)**.

| Predictor | Estimate | SE | t value | P-value |
| --- | --- | --- | --- | --- |
| Intercept (Neg Binomial) | 4.61 | 0.51 | 9.09 | < 2e-16 |
| Length of time series | -0.13 | 0.044 | -2.90 | 0.0038 |
| Always absent | 1.68 | 0.72 | 2.35 | 0.019 |
| Historical | 0.29 | 0.72 | 0.41 | 0.68 |
| Mean | 1.39 | 0.72 | 1.94 | 0.053 |
| Pooled mean | 1.61 | 0.72 | 2.25 | 0.025 |
| Uniform | -0.52 | 0.72 | -0.73 | 0.47 |
| Length*Always absent | 0.0070 | 0.062 | 0.11 | 0.91 |
| Length*Historical | -0.021 | 0.062 | -0.35 | 0.73 |
| Length*Mean | 0.0076 | 0.062 | 0.12 | 0.90 |
| Length*Pooled mean<sup>1</sup> | 0.090 | 0.062 | 1.46 | 0.15 |
| Length*Uniform | 0.27 | 0.062 | 4.31 | 1.86E-05 |

<sup>1</sup> If the analysis is releveled to the Pooled Mean model, the length of time series variable is not significant for this model ( $p = 0.40$ ).

## Footnotes

1 Model predictions would differ if a pooled mean incidence null were used. However, the Pooled Incidence model represents a more sophisticated version of that null model, hence we did not evaluate a non-probabilistic pooled mean incidence model.

2 Using the MASS package in R (Venables & Ripley 2002).

3 Using the arima function in R (Ripley 2002).

